# Vascular inflammation exposes perivascular cells to SARS-CoV-2 infection

**DOI:** 10.1101/2022.04.05.487240

**Authors:** Cristiane Miranda Franca, Amin Mansoorifar, Avathamsa Athirasala, Ramesh Subbiah, Anthony Tahayeri, Luiz Bertassoni

## Abstract

Pericytes stabilize blood vessels and promote vascular barrier function. However, vessels subjected to pro-inflammatory conditions have impaired barrier function, which has been suggested to potentially expose perivascular cells to SARS-CoV-2. To test this hypothesis, we engineered pericyte-supported vascular capillaries on-a-chip, and determined that the extravasation and binding of spike protein (S1) on perivascular cells of inflamed vessels to be significantly higher that in healthy controls, indicating a potential target to understand COVID-19 vascular complications.

## Introduction

The coronavirus disease-2019 (COVID-19), caused by the betacoronavirus SARS-CoV-2, has remained a serious public health emergency that has resulted in an enormous global disease burden^1^. COVID-19 primarily presents with respiratory symptoms including cough, shortness of breath, and fever. In more severe cases, the disease can progress to acute respiratory distress syndrome, respiratory failure, cytokine storm, and fatal multiple organ failure^1^. Worse prognosis and higher death rates are seen in patients with co-morbidities associated with chronic inflammatory conditions, such as heart disease, diabetes, asthma and cancer, in which endothelial dysfunction is known to be a key determinant ^2-4^.

Blood vessels are formed by a thin layer of endothelial cells surrounded by perivascular cells, identified as mural cells, or vascular smooth cells for larger vessels and pericytes for capillaries, venules, and arterioles ^5-7^. Perivascular cells contribute to the biological stability of vessels, by orchestrating paracrine signalling for vessel maintenance ^6, 8^, regulating vascular tone ^9^, controlling permeability ^10^, modulating coagulation ^11^ and immunologic responses ^12^.

Both endothelial cells and pericytes express angiotensin-converting 2 enzyme receptor (ACE2), which is the main binding site for SARS-CoV-2 spike protein subunit 1 (S1) ^13,14,6^. Despite the large body of evidence on the vascular complications of COVID-19, and recent reports on the contribution of pericytes to the evolution of the disease, the implications of perivascular cells and inflammation to SARS-CoV-2 infection have remained highly ellusive.

Recently Wang et al., demonstrated that pericytes can become ‘replication hubs’ for SARS-CoV-2 in the brain, mediating COVID-19-related neuroinflammation ^15^. Also, a reduction in the vascular coverage by pericytes has been documented in the heart and lung of COVID-19 patients suggesting that SARS-CoV-2 may affect the microvasculature specifically targeting pericytes ^16, 17^. Those studies shed light on that for specific organs, there is a potential preference of SARS-CoV-2 for binding to perivascular cells rather than endothelial cells, but whether a previous disruption of the vascular barrier can dictate if viral particles will bind differently to endothelial cells or perivascular cells, remains to be addressed.

Here we hypothesize that in healthy vascular capillaries with preserved barrier function, SARS-CoV-2 should have restricted access to perivascular cells, thus causing minimal vascular damage and controlled immune activation. On the other hand, vessels with altered barrier function, such as that observed in the presence of pro-inflammatory mediators like TNFα^10^, viral vascular extravasation should be elevated, thus potentially establishing an important link between inflammation, perivascular cells and vascular function ^2, 10, 18^. This relationship would shed new light onto the pathogenesis of COVID-19, and help explain the widespread implications of the disease in various vascularized tissues and organs in the body.

To test this hypothesis, we used two well-established on-chip models of vascular capillaries. We then enhanced these models by engineering endothelial capillaries that are supported with mesenchymal stromal cell (MSC)-derived perivascular cells/pericytes, and sought to understand the effect of vascular inflammation on the preferential binding of S1 protein in perivascular cells. Our results suggest inflammation mediated by TNFα significantly increases the access of the SARS-CoV-2 S1 spike protein to perivascular regions, thus helping explain the exacerbated outcomes of COVID-19 on patients with inflammatory co-morbidities.

## Results and discussion

A previously published microfluidic device (AIM Biotech) constituted of a 1.3-mm-wide central hydrogel region flanked by two lateral media channels^19, 20^, was used to engineer a microphysiologic model of perivascular cell-supported microvasculature on-chip. We encapsulated a cell suspension of GFP-expressing human umbilical vein endothelial cells (GFP-HUVECs) and human mesenchymal stromal cells (hMSCs), in a 4:1 HUVEC:hMSC ratio, in a collagen-fibrin hydrogel and seeded in the central chamber of the devices. Devices were cultured with EGM media supplemented with 50 ng/ml of vascular endothelial growth factor (VEGF) and 100 ng/ml angiopoietin 1 (Ang-1) under bidirectional agitation on a 2D rocker. After 48 h, a perivascular cell-supported network of capillaries was formed (**Figure 2A**).

**Figure 1.**
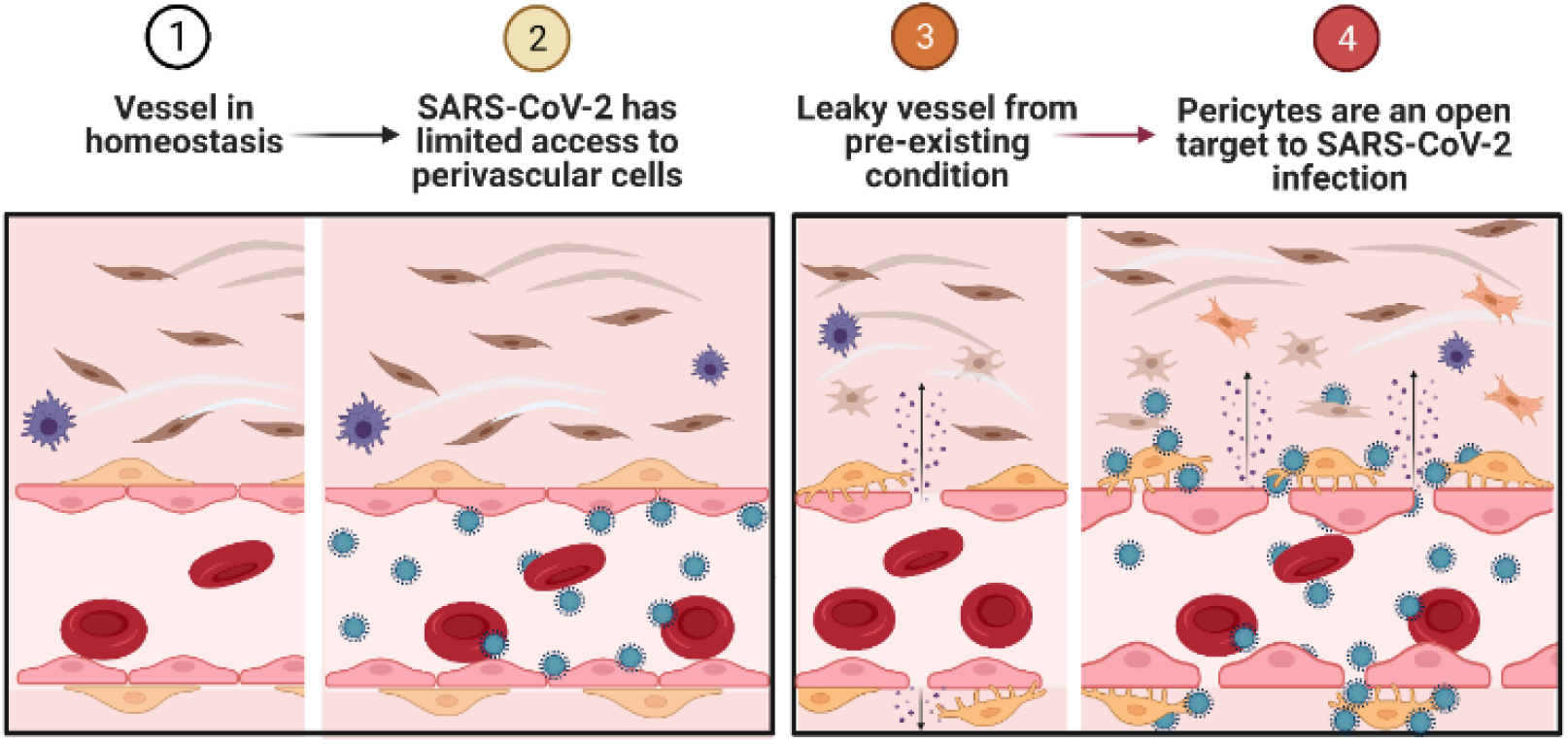
We hypothesize that inflammatory conditions that lead to disruption of vessel barrier function, may expose perivascular cells to viral particles that are circulating in the blood stream, amplifying the vascular injury related to COVID-19, partially explaining vascular complications of the disease.

**Figure 2.**
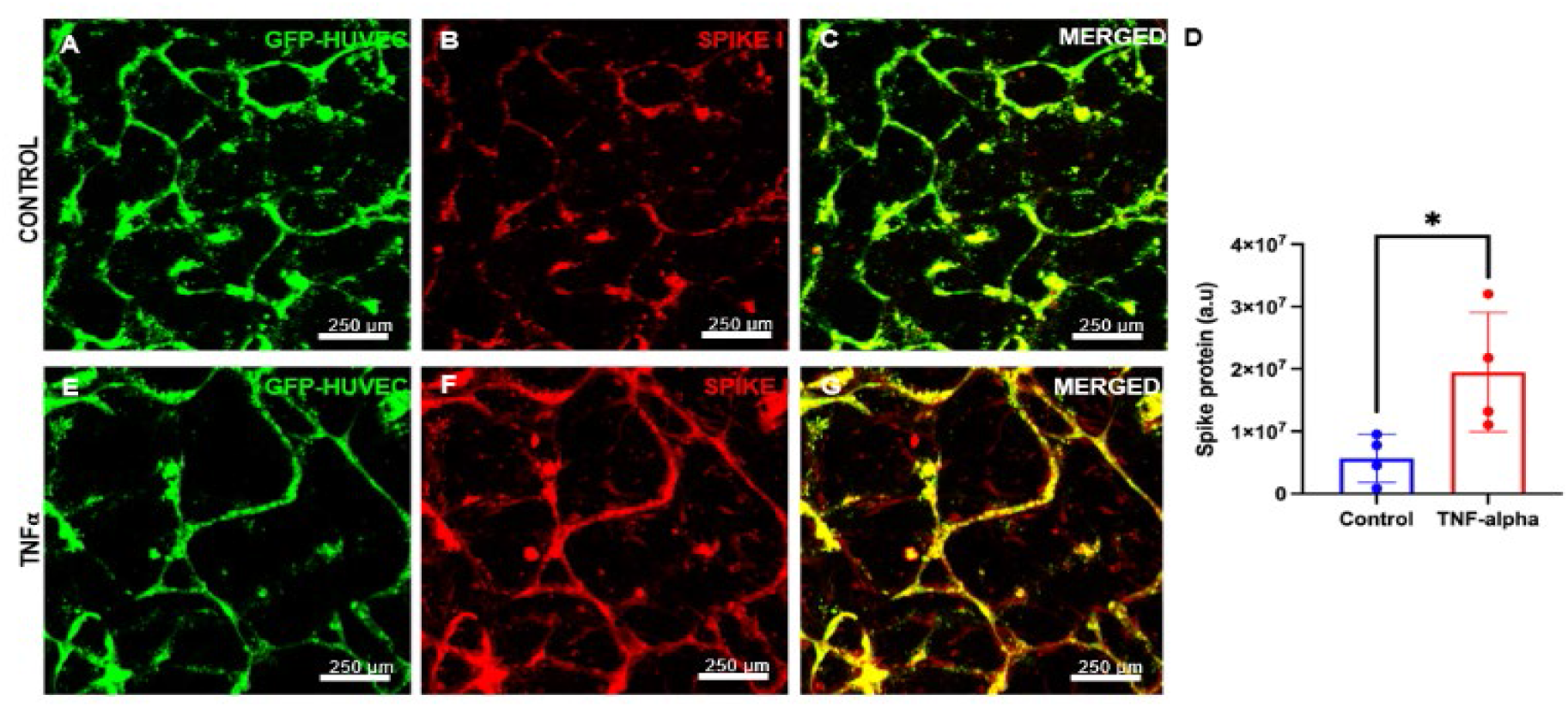
Vessels in homeostasis show binding of S1 protein concentrated in junctions and the intravascular space (A-C). Vessels previously exposed to TNFα have an augmented S1 binding protein both in the intra and extravascular space. (Student’s t test, p<0.05)

Vascular destabilization and inflammation was accomplished by treating vessels with TNFα, a proinflammatory cytokine that can cause loss of endothelial intercellular junctions ^10^. To compare the binding of spike protein on homeostatic and inflamed vessels, both devices were then incubated with spike I protein (S1) (SinoBiological, 40589-V08B1) (300 ng/mL) for 15 min according to manufacturer’s protocol. Vessels were fixed, stained, and visualized with a confocal microscope to determine the localization of the S1 protein on the vessels and perivascular area. For experimental details, see **Supplementary Information**.

S1 bound to both treated samples and controls (**Figure 2**), but were significantly more attached to the capillaries that were treated with TNFα (**Figure 2 D-G**). TNFα-treated capillaries were characterized by an even distribution of the spike protein continuously co-localized with the microvascular network also with cells within the extracellular matrix at the perivascular area (**Figure 2 F,G**). Conversely, control vessels showed an irregular distribution of the spike protein in the capillaries, more concentrated in the junction areas. Few spots of spike protein were observed in the perivascular space of control vessels (**Figure 2 B,C**).

We then engineered perivascular cells-supported single vessels (160 μm) using a previously published protocol ^21^. Briefly, an acupuncture needle was inserted in a PDMS mold with a central chamber, which was filled with type 1 collagen. After collagen assembly, the needle was removed leaving a channel that was seeded first with hMSCs and later with GFP-HUVECS. Within 48 hours the barrier function of the perivascular cells-supported 3D vessel was established ^10^. We used the same concentration of TNFα (50 ng/mL) that has been reported to impair vascular barrier function in these vessels, and loaded the SARS-CoV-2 spike protein as described above. Vessels were fixed, stained, and imaged to quantify whether S1 protein would bind preferentially to endothelial cells or pericytes under inflammatory conditions. Control vessels presented a thin homogeneous endothelialized layer of ECs co-localized with perivascular cells displaying a dim stain of spike protein bound to the vessel wall and in some migrating cells within the adjacent connective tissue (**Figure 3A**).

**Figure 3.**
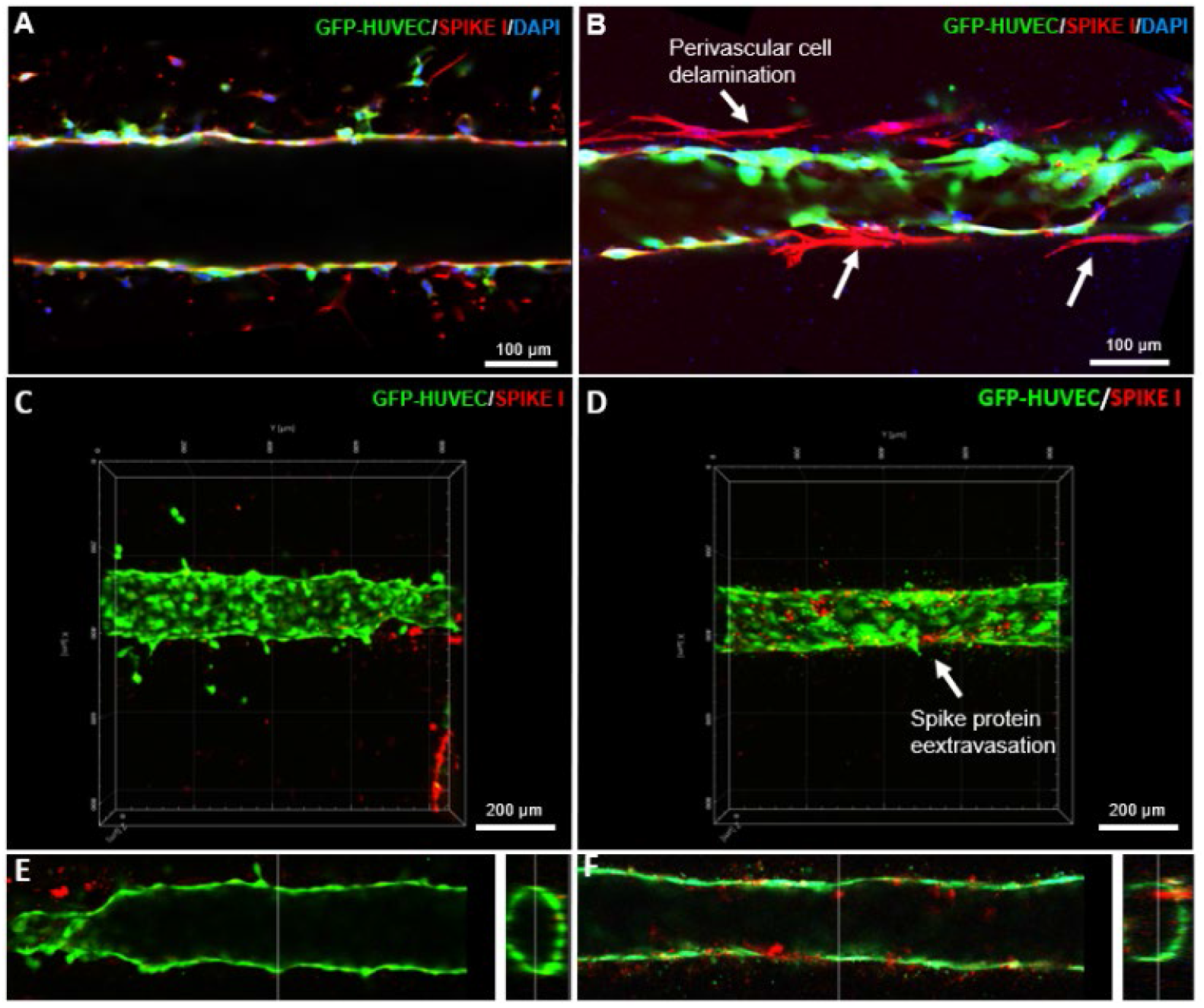
Blood vessels supported by perivascular cells in a state of homeostasis (A) show the spatial correlation of perivascular cells and endothelial cells (GFP-HUVECs) while TNFα-treated vessels depict delamination of perivascular cells, and endothelial cell thickening (B). Diffusion of spike proteins through the vessel lumen demonstrates that spike proteins leak off TNFα-treated vessels (D.F), but not from controls (C.E).

Conversely, the TNFα-treated samples showed S1 predominantly bound to delaminating perivascular cells (**Figure 3B**). Three-dimensional reconstruction of the vessels immediately after perfusing S1 protein in the vascular lumen showed virtually no spike protein outside control vessels, while TNFα led to the loss of barrier function which allowed S1 proteins (**Figure 3D, red**) to cross the endothelial barrier remain outside the vessel (**Figure 3C**). Exposure to TNFα caused the number of migrating perivascular cells to double as compared to controls (**Figure 4A**). Moreover, perivascular cells that had contact with TNFα increased their size **(Figure 4B**), acquiring a migratory phenotype with elongated, stellate-shaped cytoplasm, major processes oriented parallel to the long vascular axis, and several pronounced filopodia directed outward the vessel, within the surrounding extracellular matrix.

**Figure 4.**
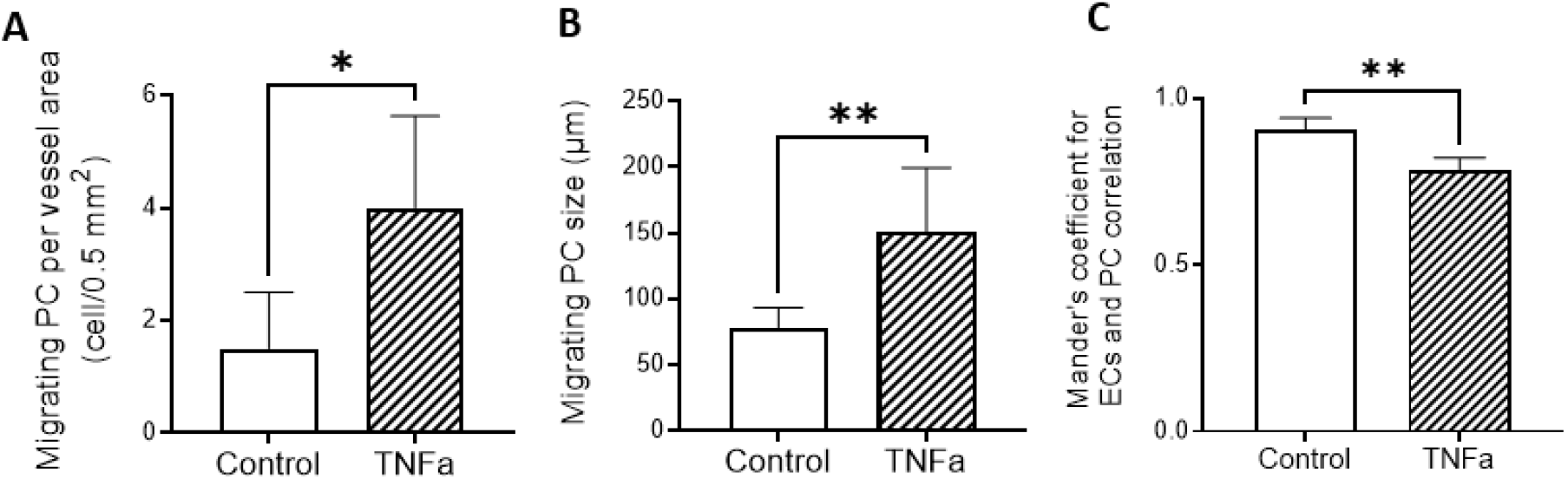
TNFα treatment induced perivascular cells to acquire a migratory phenotype (G), increase in size (H), and decreased the spatial correlation between those cells and ECs (I). One-way ANOVA, post hoc Tukey, α= 0.05.

Next, we investigated whether S1 protein would change from an intraluminal location to a more perivascular site in inflamed vessels. To that end, we normalized the background intensity of the samples and used Mander’s colocalization coefficient to evaluate co-occurrence, of S1 protein and GFP, in controls and TNFα treated vessels. This coefficient ranges from 0 to 1, which represents no overlap and complete overlap of signals respectively ^22, 23^. Control samples presented a spatial correspondence of 0.9 (0.86±0.03) while TNFα-treated vessel had a lower spatial correlation between GFP and S1 signal (0.78±0.3) indicating that S1 protein distribution is spatially broader in inflamed vessels.

Understanding the vascular mechanisms associated with perivascular cells in SARS-CoV-2 infection is crucial to providing more efficient therapeutic approaches ^24, 25^. In a cohort of 44,000 people with COVID-19 in China, three main different clinical presentations were observed in the patients: 81% showed mild symptoms, 14% had severe symptoms and 5% developed respiratory failure associated or not with multi-organ dysfunction. The overall worldwide case fatality ratio has ranged from 1.8 to 2.3%, so the varied clinical presentation, despite a nearly identical viral genome during the early phases of the pandemic, suggests a significant host influence on the clinical manifestation ^24^. In our study, we show a proof-of-principle evidence that vessels with a compromised barrier function mimicking a baseline inflammatory condition, exposed perivascular cells to viral proteins, which can potentially be one of the causes of the multisystemic clinical features of COVID-19 in patients with pre-existing inflammatory diseases.

There are several challenges related to perivascular cells that have hindered the study of perivascular cells-related vascular mechanisms in COVID-19 ^26^. For example, the morphological localization of perivascular cells sharing the basement membrane with endothelial cells and the lack of exclusive markers for reproducible histological identification of perivascular cells are obstacles for prompt identification of those cells using regular tools of investigation ^26^. Mature perivascular cells express αSMA, but this marker may reflect not only perivascular cells but also parenchymal cells and mesothelial derived myofibroblasts, which are implicated in tissue repair ^5^. In this case, microfluidic devices designed to engineer perivascular cells-supported vasculature disclose enormous possibilities to understand the actual role of perivascular cells and perivascular cells in the onset and amplification of COVID-19 vascular manifestations and how underlying conditions, such as inflammation and diabetes can contribute to vascular damage associated with COVID-19.

## Conclusions

In conclusion, we used vasculature-on-a-chip model systems to present a proof-of-principle evidence that inflamed vessels may no longer offer a barrier to viral spread from the primary viral infection site, consequently exposing perivascular cells to more abundant viral loads, and potentially amplifying the vascular injury related to COVID-19. This data suggest that perivascular cells could be a potential target to understand COVID-19 vascular complications.

## Author Contributions

CMF – data curation, formal analysis, methodology, validation, visualization, writing; AM – data curation, investigation, methodology, validation (writing – original draft); AA – formal analysis, investigation, methodology, writing (original draft) ; RS – data curation, methodology, writing (original draft); AT – methodology; LEB -conceptualization, funding acquisition, supervision, writing (review & editing).

## Acknowledgments

This project was supported by funding from the National Institute of Dental and Craniofacial Research (R01DE026170, 3R01DE026170-03S1 and 3R01DE026170-03S2 to LEB), and 1K01DE030484-01A1 to CMF.

## Experimental Section

### Cell culture

Human umbilical vein endothelial cells expressing green fluorescent protein (GFP-HUVECs) (C2519A, Lonza, Basel, Switzerland) were cultured in a supplemented (EGM-2 bullet kit, LONZA) endothelial cell growth medium (Lifeline Cell Technology, California, USA). Human bone marrow mesenchymal stem cells (hMSCs, RoosterBio, Maryland, USA) were cultured in alpha Minimal Eagle’s medium (α-MEM) (Gibco, Carlsbad, USA) with 10% FBS and 1% penicillin/streptomycin. Cell media was changed every other day and cells were passaged when reaching a confluency of 80–90%. HUVECS from passages 4–6 and hMSCs from passages 2–4 were used for all the experiments. All cells were maintained in a humidified incubator (5% CO2, 37oC).

### Fabrication of devices for microvasculature

We used commercially available microfluidic devices (AIM Biotech 3D cell culture chips – cat # DAX01, Sigma Aldrich) to engineer the microvasculature. Each device contains a set of 3 chips constituted by a 1.3-mm-wide central hydrogel region flanked by two lateral media channels, as previously described ^19, 20^. To prepare cell-laden collagen-fibrin hydrogels, acid solubilized Type 1 collagen from rat tail tendon (3 mg/mL, BD Biosciences) was reconstituted in an ice bath to a final concentration of 2.5 mg/mL, and the pH was adjusted to 7.4 by neutralizing the hydrogel precursors with 1N NaOH, and a cell suspension of HUVECs and hMSCs in EGM, in a 4:1 HUVEC:MSC ratio, 18 × 10^6^ cells/mL was added to the collagen. Next, 20 μl aliquots of cell-laden collagen were mixed with 10 μl of a thrombin solution (8 U/mL) and added to 10 μl of 5 mg/mL fibrinogen solution (cat #f3879, Sigma). A final concentration of 9×10^6^ cells/mL was used in this study. Each microfluidic device was placed for 15 minutes in a humidity chamber under the hood, then transferred to the 37°C incubator for 30 minutes for complete polymerization. Subsequently, the cell-laden hydrogel was cultured with EGM media supplemented with 50 ng/ml VEGF and 100 ng/ml Ang-1 under bidirectional agitation on a 2D rocker to induce interstitial flow during culture.

### Fabrication of microfluidic devices for single channel studies

A CAD program was used to design a mold comprised of two reservoirs connected by a central channel and a chamber. We then used a 3D printer (CADworks3D μMicrofluidic printer) and resin (Master Mold resin, CADworks) to print the positive molds. Printed μmolds were cleaned in methanol in three rinses of 2 min each under agitation, subsequently were cast with polydimethylsiloxane (PDMS, Sylgard 184, Dow-Corning), and left to cure overnight at 80°C, as previously described ^10, 21, 27^. Next, PDMS was peeled off from the resin mold, two reservoirs were prepared using a 5-mm biopsy punch for the media reservoirs and 1-mm biopsy punches for the collagen ports. Molds were cleaned with ethanol, and immediately plasma bonded to a glass coverslip. Assembled devices were autoclaved, then treated with 1% (w/v) glutaraldehyde (Sigma) for 15 min, rinsed three times with distilled water (DIW), and left overnight in DI water to remove any trace of glutaraldehyde. To mold cylindrical channels, sterile 160-μm-diameter acupuncture needles were immersed in 0.1% BSA solution for at least 30 min, then inserted into the central channel of the devices approximately 200 μm above the glass coverslip surface. Rat tail collagen type I (3.0 mg/mL, BD Biosciences) was prepared according to the manufacturer’s protocol, pipetted into the middle chamber, and allowed to polymerize at 37oC, 5% CO_2_, for 30 min. The reservoirs were then filled with EGM and devices were returned to the incubator for additional 4 hours. Afterward, the needles were carefully removed with a pair of tweezers, and the cell medium was replaced by a fresh medium, and the devices were left in the medium overnight to rinse and equilibrate the gel at 37°C, 5% CO_2_. Next, a suspension of 10^6^ hMSC/mL was seeded into the devices. The cells were allowed to adhere to the top surface of the channel for 5 min and then we performed a second seeding, flipped the device to allow the cells to adhere to the channel bottom. As soon as the channels reached the appropriate cell density, the cell suspension was removed and replaced by a fresh medium, and left incubating for 3-4 hours. HUVECs (4 × 10^6^ cells/mL) were seeded following the method described for the hMSCs. After cell attachment, EGM medium was added into the reservoirs, and devices were placed on a platform rocker (BenchRocker) (100% humidity, 37°C, 5% CO_2_). For a detailed protocol see ^10, 21^. After 48 h the vascular channels were ready to perform the assays with TNFα and Spike I proteins as described below.

### Treatment with TNFα and spike protein binding assay

To test the hypothesis that inflamed vessels enable more spike protein to bind perivascular cells, after 3 days, the formed vessels were treated with TNFα (50 ng/mL) for 1 h, were rinsed with fresh EGM, then incubated with spike I protein (SinoBiological, 40589-V08B1) (300 ng/mL) for 15 min according to manufacturer’s protocol. According to the type of device, the solutions were added to the 5-mm reservoirs (if single channel) or into the lateral channels (if the AMI biotech device). Devices were rinsed with DPBS and fixed with formalin 10% (v/v, Sigma) formaldehyde for 30 min under agitation on a bench rocker.

#### Immunofluorescence and imaging

For both types of devices, samples were fixed and permeabilized using 0.1% (v/v) Triton X-100 for 15 minutes. Next, samples were blocked with 1.5% (w/v) bovine serum albumin (BSA, Sigma Aldrich) in DPBS for 1 h, treated with Image-iT FX signal enhancer (Invitrogen, CA) for 30 min. For detection of α SMA expression, samples were incubated with mouse monoclonal antibody against α SMA (0.4 µg/ml) (MA5-11547, Thermo Fisher Scientific) (and) antibody against ACE2 (2 µg/ml) (PA5-20040, Thermo Fisher Scientific) overnight at 4°C followed by secondary antibodies (1:200 dilution) of Goat anti Mouse IgG Alexa Flour 647 and Goat anti Rabbit IgG Alexa Flour 555 (Thermo Fisher Scientific) for 2 hours at room temperature. To visualize the localization of Spike 1 protein conjugated with Mouse IgG, samples were treated with secondary antibodies Goat anti Mouse IgG (ThermoFisher Scientific, 1:200) for 2 hours. All samples were counter stained with DAPI for 15 minutes. The 3D z-stack images were obtained in a laser scanning confocal microscope (Zeiss, LSM 880, Germany). Z-stacks were converted into TIFF files using Zen Black (Zeiss) or Imaris software (v9.1, Bitplane, Oxford Instruments, Zurich, Switzerland).

### Image analysis and quantification of perivascular cells detachment and spike protein I

The total area of vascularized tissue was automatically quantified by ImageJ, the number and size of migrating perivascular cells, was determined using ImageJ and plugins ‘cell counter’ and ‘measure’, respectively ^10^. Spike protein binding to vascular cells was quantified using FIJI image analysis software for 3 ROIs per sample (n=4). For this, the regions occupied by vasculature were identified in maximum intensity projections of each 3D image file. A mask of the boundaries of these regions were then transferred to corresponding image with the spike 1 protein. The amount of spike 1 protein bound to vascular cells in each sample was determined by the average of the intensity of spike protein (red) closely associated with the boundaries created by the mask over 3 ROIs.

## Statistical analysis

All experiments were done at least in quadruplicate. Data are presented as mean ± SD, statistical analyses were performed using GraphPad Prism (version 9, GraphPad Software, LLC) using one-way ANOVA and Tukey’s post hoc tests (α=0.05). Statistically significant difference was determined as *(p < 0.05) and **(p < 0.01), respectively.

